# Investigating structure function relationships in the NOTCH family through large-scale somatic DNA sequencing studies

**DOI:** 10.1101/2020.03.31.018325

**Authors:** Michael W J Hall, David Shorthouse, Philip H Jones, Benjamin A Hall

**Affiliations:** Wellcome Trust Sanger Institute, Hinxton, CB10 1SA, UK; MRC Cancer Unit, University of Cambridge, Hutchison-MRC Research Centre, Box 197, Cambridge Biomedical Campus, Cambridge, CB2 0XZ, UK

## Abstract

The recent development of highly sensitive DNA sequencing techniques has detected large numbers of missense mutations of genes, including *NOTCH1* and 2, in ageing normal tissues. Driver mutations persist and propagate in the tissue through a selective advantage over both wild-type cells and alternative mutations. This process of selection can be considered as a large scale, in vivo screen for mutations that increase clone fitness. It follows that the specific missense mutations that are observed in individual genes may offer us insights into the structure-function relationships. Here we show that the positively selected missense mutations in *NOTCH1* and *NOTCH2* in human oesophageal epithelium cause inactivation predominantly through protein misfolding. Once these mutations are excluded, we further find statistically significant evidence for selection at the ligand binding interface and calcium binding sites. In this, we observe stronger evidence of selection at the ligand interface on EGF12 over EGF11, suggesting that in this tissue EGF12 may play a more important role in ligand interaction. Finally, we show how a mutation hotspot in the NOTCH1 transmembrane helix arises through the intersection of both a high mutation rate and residue conservation. Together these insights offer a route to understanding the mechanism of protein function through *in vivo* mutant selection.

## Introduction

Mutational scanning experiments exhaustively mutate a protein or protein domain and measure the phenotypic consequences of each mutation (1). They can provide information both about the effects of individual mutations and about protein function and structure (1). However, these experiments are carried out *in vitro (1),* an environment which can substantially alter cell phenotype (2, 3). Over the last decade, DNA sequencing has enabled the detection of vast quantities of mutations occurring in normal tissue samples (4–8). These may be thought of as large-scale, *in vivo,* mutational phenotype assays. *NOTCH1* is one of the most frequently mutated genes in these studies, which have provided many more mutations than previously identified in large scale cancer genome and developmental mutation studies (9–12).

Mutations are acquired by cells as a result of mutagen exposure (e.g. tobacco, alcohol or ultraviolet light) or cell intrinsic processes (13). The pattern of mutations acquired, known as the mutational spectrum, will depend on the mutational processes involved (13). The chance of acquiring a particular single nucleotide substitution appears to depend (at least partially) on the adjacent nucleotides (13). Many of the mutations do not alter cell behaviour and are lost through neutral drift, the process of random cell births and loss in which by chance some cell lineages expand and others are lost (14). Positively selected (driver) mutations confer a proliferative advantage to the mutant cell, thus increasing the number of descendant cells that inherit the mutation (15), and are therefore more likely to generate a clone large enough to be detected by DNA sequencing (16).

To detect genes under selection, it is assumed that mutations are generated according to the mutational spectrum (Fig. 1). In non-selected, neutral genes, where no mutations of any kind convey a growth advantage/disadvantage, the mutations detected by DNA sequencing will simply be an unbiased sample of the mutations produced by the spectrum. In genes under selection, it is assumed that certain types of mutations (missense/nonsense mutations (16), or mutations estimated by machine-learned approaches to have high functional impact (17, 18)) are more likely to alter cell phenotype than others (silent mutations, mutations with low estimated functional impact). Therefore, a positively selected gene would have more high-impact mutations detected than expected under the neutral null hypothesis (16–18) (Fig. 1). Conversely, negative selection in a gene would lead to fewer detected high-impact mutations than expected under the null hypothesis.

**Figure 1:**
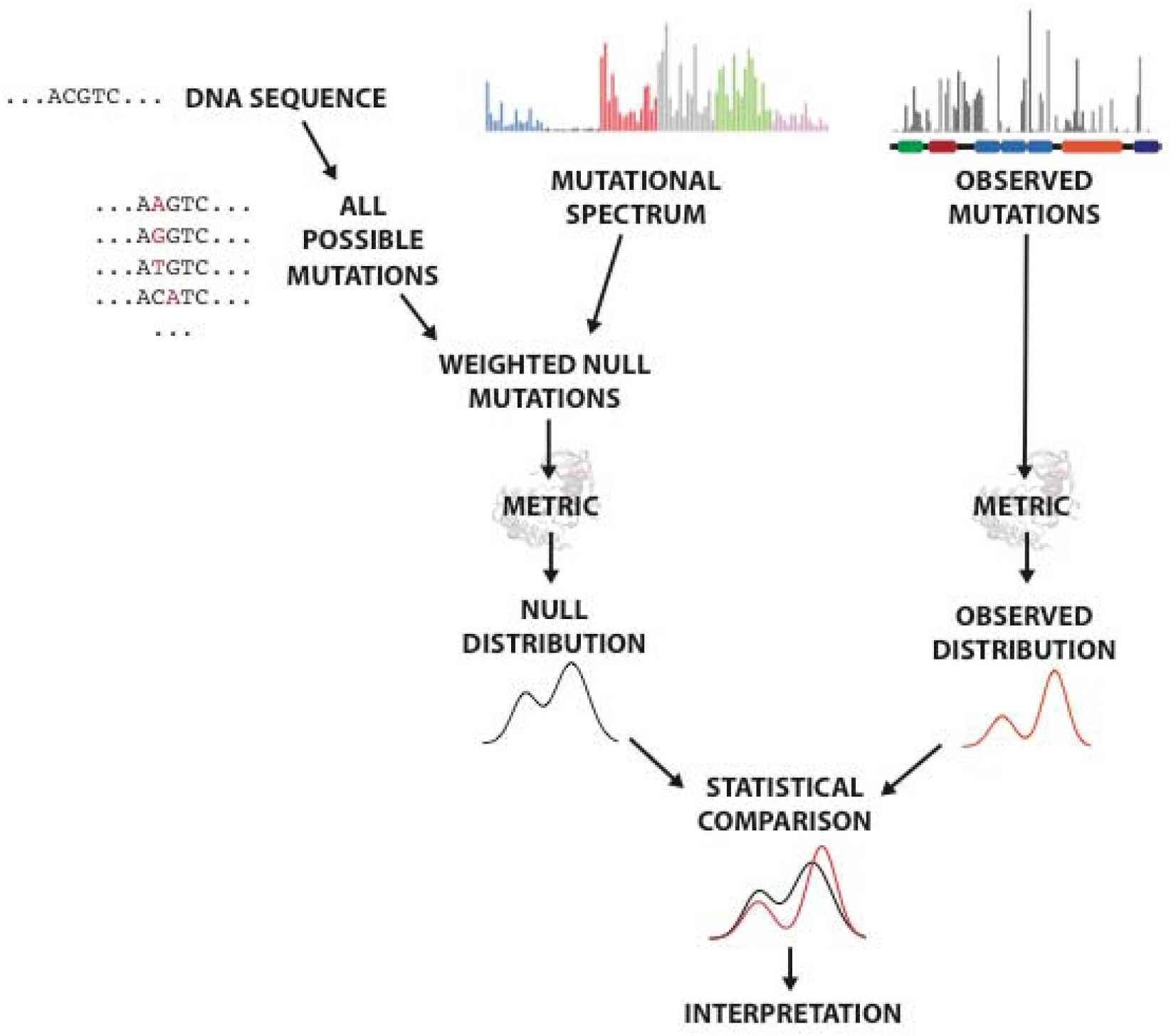
Schematic of the statistical test for a selected feature. The null hypothesis assumes that the mutations appear based on the mutational spectrum, and there is no selection or other bias occurring. The alternative hypothesis is that some selection occurs, and that it correlates with the tested metric. For example, in this study the metrics used were ΔΔG, and whether the mutation was within/not within a specified set of residues (scored 1/0) **(methods).** Any metric can be used if it can be described at least semi-quantitatively for some mutations in the region and may correlate with selection. To generate the metric distribution under the null, all possible single nucleotide mutations were generated for the gene/region to be tested, weighted by the mutational spectrum and scored using the metric. This was then compared to the distribution of metric scores for the observed mutations using a statistical test **(methods).** The comparison determines if the mutations with a high/low metric score are more strongly selected than the rest of the mutations inthe tested region, and therefore must be interpreted while considering other causes of selection in the region **(Supplementary text-Testing Region).**

In this study, we use the same principle to look in more detail at patterns of selection within single genes **(Fig. 1).** In a similar manner to mutational scanning experiments, by identifying common features of selected mutations we can gain information about how the wild type and mutant proteins are functioning. Here we focus on *NOTCH1* and *NOTCH2,* membrane bound cell surface receptors **(Fig. 2a)** in a pathway that regulates cell fate (19). These genes are critical regulators of normal cellular function in development and adult tissues and are mutated to activate or block function in difference cancers (20, 21). The extracellular domains of NOTCH genes contain up to 36 epidermal growth factor (EGF) repeats **(Fig. 2b,c).** NOTCH ligands, from the Delta-like and Jagged families (22), expressed by adjacent cells and bind to a subset of the EGF repeats (19) **(Fig. 2a).** This binding triggers a cascade of cleavage events, resulting in the cleavage of the NOTCH transmembrane helix and the release of the NOTCH intracellular domain (NICD), which travels to the nucleus and forms part of a transcription factor complex increasing the expression of NOTCH target genes (19) **(Fig. 2a).**

**Figure 2:**
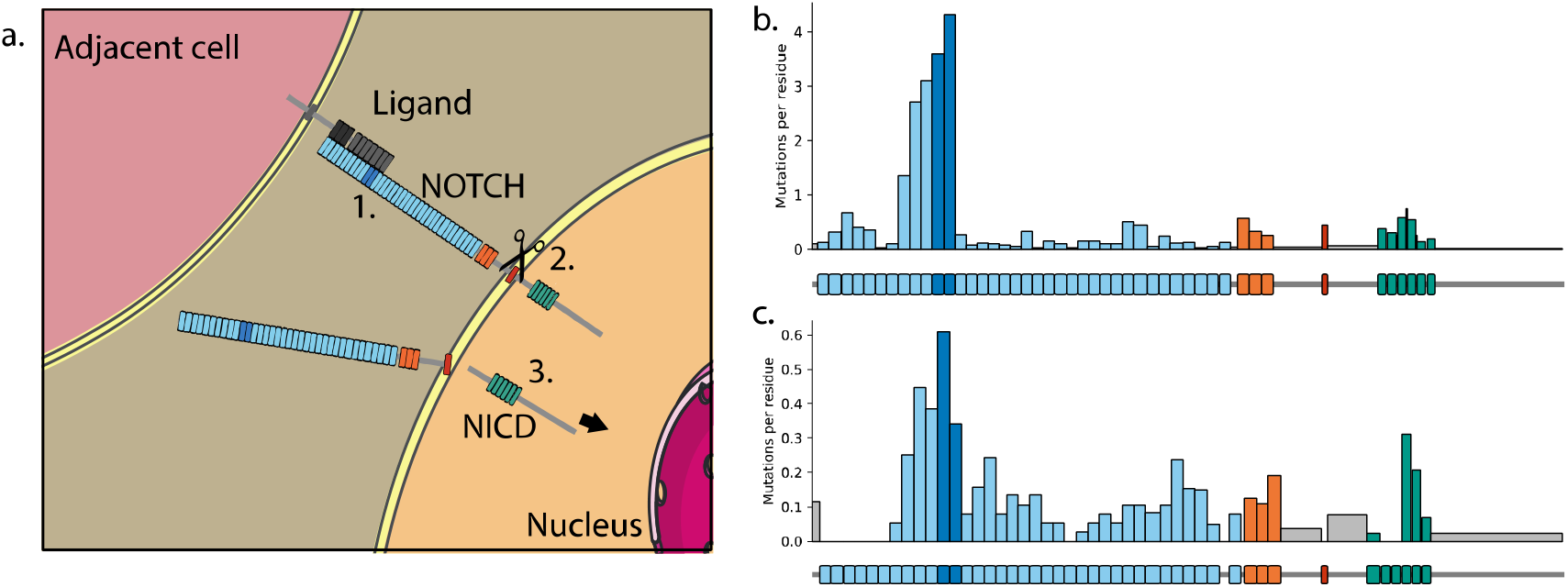
NOTCH activation, structure and mutation, **a.** (1) NOTCH is activated when a ligand from an adjacent cell binds with some of the NOTCH EGF repeats (blue, EGF11-12 shown in dark blue). (2) Ligand binding triggers a series of cleavage events, resulting in cleavage of the NOTCH transmembrane region (red) by ysecretase. (3) This releases the intracellular domain of NOTCH (NICD) which travels to the nucleus to form part of a transcription factor, **b-c.** Missense mutation frequency across the domains of NOTCH1 **(b)** and NOTCH2 **(c)** (60) in normal human oesophagus (4). Where the gap between domains is only a single residue, mutations from this residue are included in the subsequent domain. EGF repeats, blue; EGF11-12, dark blue; LNR repeats, orange; transmembrane region, red; ankyrin repeats, green; other regions, grey.

Structural studies have revealed that EGF11-12 are particularly crucial for ligand binding (19, 23), though EGF repeats 8-10 are also known to be involved in ligand interaction (24, 25). EGF11 and EGF12 of NOTCH genes bind to the Delta–Serrate–Lag-2 (DSL) and C2 domains of the ligand respectively (26–28). Based on the conservation of interface residues in the four NOTCH genes and the NOTCH ligands, it has been proposed that the more conserved EGF11-DSL interface acts as a consistent binding point for ligand binding, whereas the EGF12-C2 interface may determine which ligand-notch pairs can bind (26), although it has also been argued that EGF12 contains the major binding site for the ligand Delta-like 1 (Dll1) (29). Many of the EGF repeats bind to calcium ions, which add rigidity to the structure, help fix the relative orientation of adjacent EGF repeats (30), and are required for ligand binding (29). A calcium binding site in EGF12 in particular has been found to be critical for Notch-Dll1 binding (29).

The NICD is released by cleavage of the transmembrane helix of Notch at the S3 site by the multiprotein γ-secretase complex (31). The exact site of cleavage can vary slightly, but the S3-V cleavage (between G1753 and V1754 in *NOTCH1)* produces the most stable form of NICD due to N-end rule degradation (32) and therefore is likely responsible for the majority of Notch signalling (19). Mutations in the vicinity of the S3 cleavage site of NOTCH may affect Notch signalling by reducing S3 cleavage efficiency (33) or by leading to the production of less stable NICD species (32).

NOTCH1 inactivation is a strong driver of clonal expansion in healthy skin and oesophageal epithelium (4, 6, 8). To study the structural properties of inactivating NOTCH mutations, we used a data set of approximately 7000 mutant clones detected through DNA sequencing of normal oesophageal epithelium (4). It was found that missense mutations in NOTCH1 (n=951) cluster in certain regions of the protein, with the highest concentration of mutations in the ligand-binding EGF repeats 11 and 12 **(Fig. 2b)** (4). Although less frequent, missense mutations are also scattered across other regions of the protein, including the transmembrane helix **(Fig. 2b).**

*NOTCH2* is also detected as a positively selected mutant gene in the same data set, although with fewer mutant clones (234 missense mutations) and a lower dN/dS ratio (4), indicating it is under weaker selection than *NOTCH1.* It shows a similar, though less pronounced, increase of mutation density in EGF11-12 **(Fig. 2c).**

In this study we develop a statistical methodology that shows that missense mutations in *NOTCH1* EGF11-12 that destabilise the protein, alter the ligand-binding interface or disrupt calcium-binding sites are significantly selected. We find that the same features are selected for in *NOTCH2* EGF11-12. We further show that the selection of interface mutations is stronger in EGF12 of *NOTCH1* than in EGF11, while destabilising mutations are equally selected for both EGF repeats. We also find another major hotspot mutation in the transmembrane region of *NOTCH1* which is the only mutation on a highly conserved residue that also has a high expected mutation rate, underlining the need to consider the mutational spectrum when interpreting mutation selection from sequencing studies.

The statistical method used here detects differences in the *relative* strength of selection. This means that, in regions with multiple selected features, each feature can hinder the detection of the other features. We show how this confounding effect can be reduced by constructing models for the null and alternate hypotheses that condition on known selected features while testing for another. Together, our results highlight the importance of examining both the biophysical mechanism of mutant action alongside the mutagenic processes that create them. It further provides a workflow for understanding the mechanisms of selection in genes.

## Results

### EGF repeat mutations cause misfolding and disrupt ligand and calcium binding sites

We began by investigating the selected features of missense mutations in the critical *NOTCH1* ligand-binding EGF repeats 11-12. Recurrently mutated residues in this region include cysteines in disulphide bonds, buried glycines and hydrophobic packing residues (4). These would all be expected to affect the stability of the protein (34, 35) and could prevent the structure from folding into the correct shape to bind with the ligand (26, 27). We therefore began by testing whether there is selection for destabilising mutations.

The stability of a protein is determined by the protein folding free energy, ΔG, which is the difference in Gibb’s free energy between the folded and unfolded form of a protein (36). A mutation may alter ΔG (this change is called ΔΔG) and therefore stabilise or destabilise the protein (36). We used FoldX (36) to calculate the ΔΔG for each possible single nucleotide missense mutation in *NOTCH1* EGF11-12 using the 2VJ3 structure (37) **(Fig. 3a).** We constructed a null model of “neutral” selection, which assumes that the distribution of mutations in the region will depend solely on the mutational spectrum **(Fig. 1, methods – Mutational spectrum, Monte Carlo test).** By comparing the distribution of ΔΔG values from the observed mutations with the distribution expected under the null hypothesis **(Fig. 1),** we found a significant enrichment of destabilising mutations with high ΔΔG values (p<2e^-5^, n=308, two-tailed Monte Carlo test, **methods) (Fig. 3b).** However, we can see that many observed mutations, including recurrent hotspots, do not appear to be destabilising **(Fig. 3a).** This suggests that some mutations are selected for reasons other than destabilising the protein structure.

**Figure 3:**
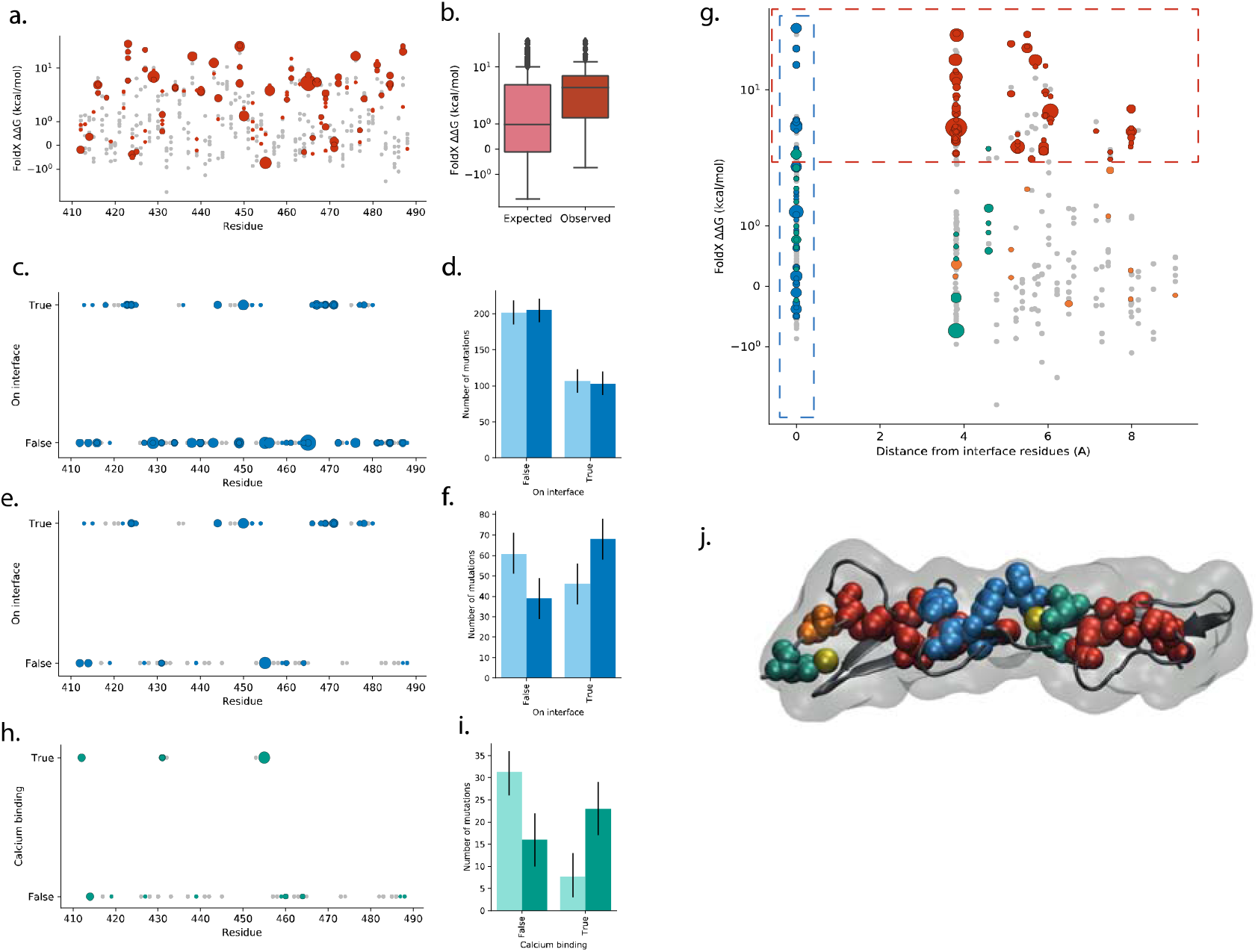
Selected features of missense mutations in *NOTCH1* EGF11-12. **a.** ΔΔG of mutations in NOTCH1 EGF11-12 calculated using structure 2VJ3 and FoldX. Single nucleotide missense mutations that occur in the human oesophagus data set, red, with marker size proportional to the number of times that mutation occurs. Single nucleotide missense mutations that do not occur in the human oesophagus data set shown in grey. **b.** Distribution of calculated ΔΔG values of missense mutations. Distribution expected under the neutral null hypothesis, light red, and the distribution observed, dark red. p<2e^-5^, n=308, two-tailed Monte Carlo test **(methods), c.** Mutations on or off the ligand-binding interface of NOTCH1 EGF11-12 **(methods).** Single nucleotide missense mutations that occur in the human oesophagus data set, blue, with marker size proportional to the number of times that mutation recurs. Single nucleotide missense mutations that do not occur in the human oesophagus data set shown in grey. **d.** Counts of NOTCH1 EGF11-12 mutations occurring on the ligand-binding interface under the neutral null hypothesis, light blue, and observed, dark blue. p=0.73, n=308, two-tailed Monte Carlo test **(methods).** Error bars show 95% confidence intervals **(methods), e.** Mutations on or off the ligand-binding interface of N0TCH1 EGF11-12 **(methods),** where mutations with a calculated ΔΔG>2 kcal/mol have been excluded. Single nucleotide missense mutations that occur in the human oesophagus data set, blue, with marker size proportional to the number of times that mutation occurs. Single nucleotide missense mutations that do not occur in the human oesophagus data set shown in grey. **f.** Counts of NOTCH1 EGF11-12 mutations with calculated ΔΔG ≤ 2 kcal/mol occurring on the ligand binding interface under the neutral null hypothesis, light blue, and observed, dark blue. p=4e^-5^, n=107, two-tailed Monte Carlo test **(methods).** Error bars show 95% confidence intervals **(methods), g.** Calculated ΔΔG plotted against distance from the NOTCH1 EGF11-12 ligand-binding interface residues calculated on the structure 2VJ3. Single nucleotide missense mutations that occur in the human oesophagus data set with marker size proportional to the number of times that mutation occurs shown in green if the residue is calcium binding, blue if the residue is on the ligand-binding interface, red if the mutation has ΔΔG>2kcal/mol, orange otherwise. Single nucleotide missense mutations that do not occur in the human oesophagus data set shown in grey. Regions containing highly destabilising mutations (ΔΔG>2kcal/mol) and mutations on the ligand-binding interface shown with dashed red and blue boxes respectively, **h.** Mutations on calcium-binding residues in NOTCH1 EGF11-12. Mutations with a calculated ΔΔG>2 kcal/mol or on a ligand-binding interface residue have been excluded. Single nucleotide missense mutations that occur in the human oesophagus data set, green, with marker size proportional to the number of times that mutation recurs. Single nucleotide missense mutations that do not occur in the human oesophagus data set shown in grey. ¡. Counts of NOTCH1 EGF11-12 mutations that are on calcium-binding residues (having excluding mutations with calculated ΔΔG > 2 kcal/mol or occurring on the ligand-binding interface), under the neutral null hypothesis, light green, and observed, dark green. p<2e^-5^, n=39, two-tailed Monte Carlo test **(methods).** Error bars show 95% confidence intervals **(methods), j.** Residues affected by recurrent mutations (occurring at least 4 times) highlighted on EGF 11-12 of structure 2VJ3. Calcium-binding residues, green; ligand-binding interface residues (which are not also calcium binding), blue; residues with highly destabilising mutations (ΔΔG >2 kcal/mol) (and which are not on the interface or calcium binding); red. D414 shown in orange, which is near to a calcium ion but not defined as a calcium-binding residue by MetalPDB. Calcium ions shown in yellow.

Another mechanism to inactivate *NOTCH1* ligand binding is disruption of the ligand-binding interface. Structures of rat *Notch1* bound to the ligands *Jag1* and *Dll4* have shown that the binding surfaces on *Notch1* are very similar for both ligands (26, 27). We used the union of both ligand binding surfaces to define the ligand interface residues ((27), **methods – Ligand-binding interface residues, Supplementary Fig. 1a, Supplementary table 1).** Under the null model, 35% of mutations were expected to appear on the interface and 33% were observed **(Fig. 3d),** and we found that the missense mutations on the interface were not significantly selected compared with the rest of EGF11-12 (p=0.73, n=308, two-tailed Monte Carlo test, **methods).** However, this does not indicate that mutations on the interface are under neutral selection, just that they are not under stronger selection than the bulk of missense mutations in EGF11-12 **(Supplementary Text – Testing region).** The presence of other mechanisms of selection could confound our analysis as they may be distal to the binding site. As we have already shown that highly destabilising mutations are positively selected, we can exclude all mutations with high ΔΔG values from both the null model and the observed data (excluding ΔΔG>2 kcal/mol, other thresholds tested in **Supplementary Fig. 1b)** and test whether, *within the non-destabilising mutations in EGF11-12,* there is an enrichment of interface mutations **(Fig. 3e).** Under this null model, 43% of non-destabilising mutations were expected to occur on the interface and 63% were observed **(Fig. 3f),** which is a highly significant increase (p=4e^-5^, n=107, two-tailed Monte Carlo test, **methods**).

After excluding misfolding mutations and those on ligand-binding sites, we find that there are still some missense mutations remaining **(Fig. 3g).** Among these, the most frequent mutation is E455K, which forms part of a calcium-binding site that is known to be crucial for ligand binding (29, 38). We used MetalPDB (38) to find the calcium-binding residues in the 2VJ3 structure **(Fig. 3g, Supplementary Fig. 1c).** Similar to the results for the interface mutations, testing with all mutations in EGF11-12 did not find a significant difference in selection (p=0.48, n=308, two-tailed Monte Carlo test, **methods, Supplementary Fig. 1d,e),** but by excluding the destabilising and interface mutations we found that the calcium-binding mutations are highly selected (p<2e^-5^, n=39, two-tailed Monte Carlo test, **methods, Fig 3h,i**).

After exclusion of misfolding, ligand-binding and calcium-binding sites, we find a single remaining frequent mutation. D414G is mutated 4 times in the oesophagus data but is not included so far in any of the categories. Visual examination of the protein however reveals that D414G is proximal to a calcium-binding site **(Fig. 3j).** We suggest that it disrupts calcium binding by reducing the affinity of the Ca^2+^ ion to the binding site by reducing the charge density around the adjacent calcium binding site. This was confirmed by Advanced Poisson-Boltzmann Solver (APBS) calculations showing that the charge surface density was substantially disrupted **(Supplementary Fig. 2).**

Among these three categories of mutational impact, we find that approximately 70% of potential single nucleotide mutations in the EGF11-12 repeats are likely to inactivate *NOTCH1* and hence be positively selected if they occur.

### Differences in selection between EGF11 and EGF12

It has been proposed that two regions of *NOTCH1* EGF11-12 may have slightly different roles in ligand interaction – Site1 on EGF12 that interacts with the C2 domain of the ligand and Site2, mostly on EGF11, that interacts with the DSL domain of the ligand (26). It may therefore be interesting to examine whether the identified categories of mutant impact are selected differently between the two key EGF repeats.

Firstly we looked at whether destabilising mutations were equally selected in EGF11 and EGF12. We constructed a null model in which the destabilising mutations were spread over all potential destabilising substitutions in EGF11-12 according to the mutational spectrum. The two-tailed alternative hypothesis was that EGF11 had either a significant increase or significantly decrease in destabilising mutations compared to EGF12. We excluded all mutations on the ligand-binding interface and calcium-binding residues from both the null model and observed mutations to remove confounding sources of selection. We found that the destabilising mutations were distributed across the two EGF repeats in accordance with

Next we looked at the ligand-binding interface residues, and split them into Site1 (EGF12) and Site2 (EGF11) (26) **(Supplementary Fig. 1a, Supplementary Table 1).** We excluded all destabilising mutations and calcium-binding residues to remove confounding sources of selection. We see significantly stronger selection of Site1 mutations compared to Site2 (p=0.04, n=53, two-tailed Monte Carlo test, **Fig. 4b).** This represents the difference in average selection of mutations in the two sites and may not characterize the overall relative importance of the two sites. However, the result may be consistent with the previous studies that have identified EGF12 as a particularly important repeat for ligand binding (29, 39).

**Figure 4:**
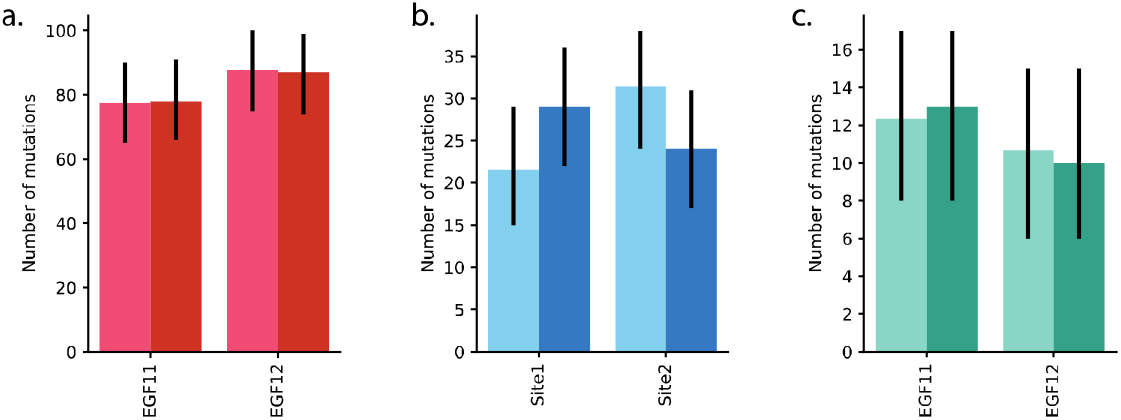
Comparison of selection in *NOTCH1* EGF11 and EGF12. **a.** Counts of NOTCH1 EGF11-12 mutations that are destabilising (with calculated ΔΔG > 2 kcal/mol), having excluded ligand-binding and calcium binding mutations, under the neutral null hypothesis, light red, and observed, dark red. p=0.95, n=165, two-tailed Monte Carlo test **(methods).** Error bars show 95% confidence intervals **(methods), b.** Counts of *NOTCH1* mutations on the two ligand binding sites (Site1 on EGF12, Site2 on EGF11, **methods),** having excluded destabilising mutations (with calculated ΔΔG > 2 kcal/mol) and calcium-binding mutations. Expected counts under the neutral null hypothesis, light blue, and counts observed, dark blue. p=0.04, n=53, two-tailed Monte Carlo test **(methods).** Error bars show 95% confidence intervals **(methods), c.** Counts of NOTCH1 EGF11-12 mutations that are on calcium-binding residues (having excluding mutations with calculated ΔΔG > 2 kcal/mol or occurring on the ligand-binding interface). Expected counts under the neutral null hypothesis, light green; counts observed, dark green. p=0.95, n=23, two-tailed Monte Carlo test **(methods).** Error bars show 95% confidence intervals **(methods)**.

There was no indication of a difference in selection of calcium binding mutations between EGF11 and EGF12 (p=0.95, n=23, two-tailed Monte Carlo test, destabilising and ligand interface mutations excluded, **Fig. 4c).**

### Selection in EGF11-12 of *NOTCH2*

Based on the results for *NOTCH1,* we considered misfolding, ligand-interface and calcium binding mutations as potentially selected groups **(Fig. 5a).** We therefore tested each category, while excluding mutations in the other two categories from both the null and observed distributions to remove confounding sources of selection. Destabilising mutations were significantly selected (p<2e^-5^, n=25, two-tailed Monte Carlo test, **methods, Fig 5b).** Both ligand-interface mutations and calcium-binding mutations were increased compared to the null models, but the trends were not quite significant (ligand interface: p=0.06, n=6, **Fig 5c;** calcium binding: p=0.053, n=5, **Fig 5d;** two-tailed Monte Carlo test, **methods).** The number of mutations exclusive to each of these two categories was very low, but we could combine the two to test whether there was significant selection of mutations that either disrupted the ligand-binding interface or a calcium-binding site (this also means we could include mutations that were simultaneously in both categories). This test revealed that there was significant selection of these features (p=0.002, n=13, two-tailed Monte Carlo test, **methods, Fig 5e).** Although the test does not conclusively show that either individual category is selected, the trends of the individual category tests and the similarities with *NOTCH1* mean we can be reasonably confident that both ligand-binding mutations and calcium-binding mutations in *NOTCH2* were under selection.

**Figure 5:**
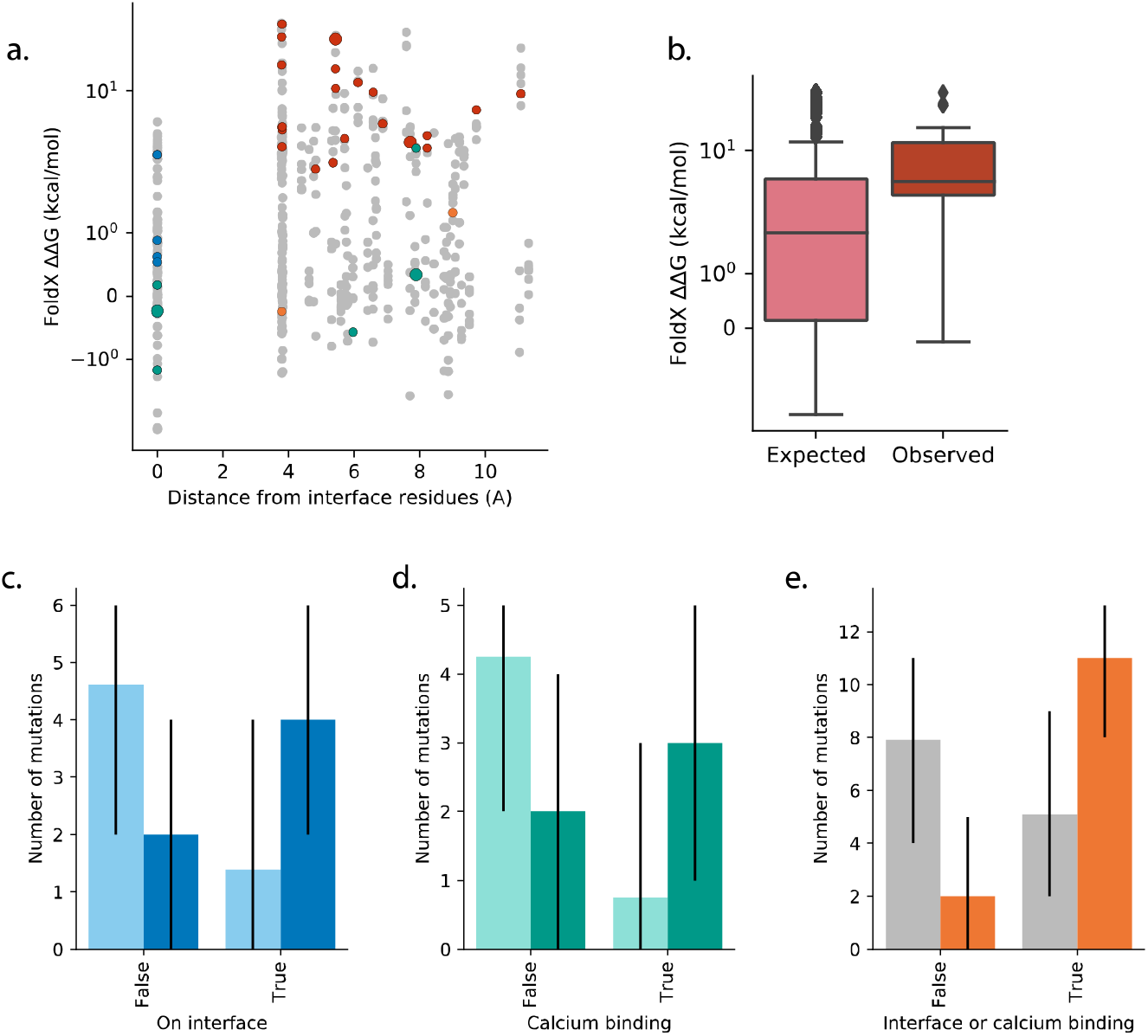
Selected features of missense mutations in *NOTCH2* EGF11-12. **a.** Calculated ΔΔG plotted against distance from the NOTCH2 EGF11-12 ligand-binding interface residues calculated on the structure 5MWB. Single nucleotide missense mutations that occur in the human oesophagus data set with marker size proportional to the number of times that mutation occurs shown in green if the residue is calcium binding, blue if the residue is on the ligand-binding interface, red if the mutation has ΔΔG>2kcal/mol, orange otherwise. Single nucleotide missense mutations that do not occur in the human oesophagus data set shown in grey. **b.** Distribution of calculated ΔΔG values of missense mutations after excluding mutations on the ligand-binding interface and calcium-binding residues. Distribution expected under the neutral null hypothesis, light red, and the distribution observed, dark red. p<2e^-5^, n=25, two-tailed Monte Carlo test **(methods), c.** Counts of *NOTCH2* EGF11-12 mutations occurring on the ligand-binding interface having excluded destabilising mutations (with calculated ΔΔG > 2 kcal/mol) and calcium-binding mutations. Expected counts under the neutral null hypothesis, light blue; counts observed, dark blue.p=0.06, rı=6, two-tailed Monte Carlo test **(methods).** Error bars show 95% confidence intervals **(methods), d.** Counts of *NOTCH2* EGF11-12 mutations occurring on the calcium-binding residues having excluded destabilising mutations (with calculated ΔΔG > 2 kcal/mol) and ligand-binding mutations. Expected counts under the neutral null hypothesis, light green; counts observed, dark green. p=0.053, n=5, two-tailed Monte Carlo test **(methods).** Error bars show 95% confidence intervals **(methods), e.** Counts of *NOTCH2* EGF11-12 mutations occurring either on the ligand-binding interface or on calcium-binding residues having excluded destabilising mutations (with calculated ΔΔG > 2 kcal/mol). Expected counts under the neutral null hypothesis, grey; counts observed, orange. p=0.002, n=13, two-tailed Monte Carlo test **(methods).** Error bars show 95% confidence intervals **(methods).**

### Hotspots in the *NOTCH1* transmembrane helix show the intersection of mutagenic processes and conservation

The EGF11-12 region was the most heavily mutated part of *NOTCH 1* in the normal human oesophagus, but there were signs of selection elsewhere in the gene. For example, we see that there is a recurrent mutation in the transmembrane region **(Fig. 2b, Fig. 6a).** In this case, we could not look for common selected features since we lacked the required large diversity of different mutations **(Supplementary Text – Correlation, causation and hotspots).** The hotspot mutation, G1753R, is on a highly conserved residue adjacent to the S3 cleavage site (40) **(Fig. 6a)** at which γ-secretase cleaves the NICD from the extracellular domain of NOTCH1. Preventing cleavage or shifting the cleavage site to produce an NICD with a shorter half-life would inactivate, or at least reduce, the *NOTCH1* signal (19, 32, 33).

**Figure 6:**
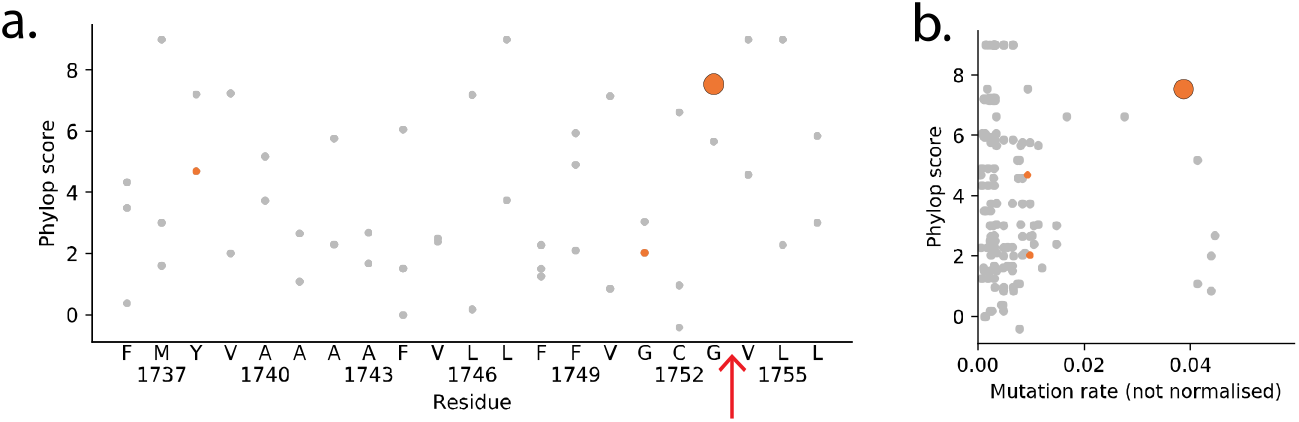
Missense mutations in the transmembrane helix of *NOTCH1.* **a.** Conservation scores from Phylop2 shown on γ-axis. Single nucleotide missense mutations that occur in the human oesophagus data set, orange, with marker size proportional to the number of times that mutation recurs. Single nucleotide missense mutations that do not occur in the human oesophagus data set shown in grey. Location of the key S3-V cleavage site indicated by a red arrow, **b.** Phylop conservation score plotted against relative expected mutation rate for missense mutations in the transmembrane helix of *NOTCH1.* Single nucleotide missense mutations that occur in the human oesophagus data set, orange, with marker size proportional to the number of times that mutation recurs. Single nucleotide missense mutations that do not occur in the human oesophagus data set shown in grey. The hotspot mutation (G1753R) is both highly conserved and has a high expected mutation rate.

Conservation of protein sequence across species is likely to be correlated with function, with residues that are more important for normal protein function more likely to be conserved (26, 28, 41, 42). G1753 is highly conserved, but so are other residues in the transmembrane region that were not mutated in the oesophagus data set **(Fig. 6a).** This possibly suggests that these non-mutated residues were not selected as strongly as G1753, despite the high conservation scores. However, looking at both the conservation score and the mutational spectrum, we see that the G1753R was the only highly conserved mutation that also had a high probability of being mutated **(Fig. 6b),** plausibly explaining why it was this mutation in particular that appeared recurrently in the data set.

This illustrates the importance of considering the mutational spectrum when running the test for selection. Within *NOTCH1* EGF11-12, unsurprisingly, mutations with a higher expected mutation rate were generally more mutated **(Supplementary Figs. 1f,g).** However, low sample size, imprecise mutation-rate estimation **(Supplementary text – Mutational Spectrum)** and potentially overlooked differences in functional impact mean that it is hard to precisely predict which individual mutations will be highly mutated **(Supplementary Fig. 1h).** This shows how it can be advantageous to test for mutation function selection using the bulk set of mutations instead of focusing on individual mutations.

## Discussion

Here we have adapted a method for cancer driver gene discovery to look for selected features of mutations in a gene. This has confirmed results from a previous analysis of recurrent hotspot mutations in normal oesophageal epithelium that identified positive selection of missense mutations in *NOTCH1* EGF11-12 that destabilise the protein, alter the ligand-binding interface or disrupt calcium-binding sites (4). In *NOTCH2* there were no mutations that occurred more than twice. However, by considering the mutations in bulk, we have shown that the same features are selected for as in *NOTCH1.* Apparent differences in selection between the ligand-binding interfaces of *NOTCH1* EGF11 and EGF12 parallel previous structural studies indicating EGF12 may be the most important ligand binding domain (29, 39). On the other hand, misfolding mutations appear to be equally selected across the two EGF repeats. Taken together, this is consistent with the expected behaviour: that interface mutations have a local effect, while the misfolding mutations have a larger disrupting effect so that the exact location of the mutation is not as crucial.

In the examples covered in this study we have looked at small regions of the gene. In principle, more information is potentially available from the distribution of mutations across the whole gene. For example, the ligand-binding EGF repeats were much more mutated than the rest of *NOTCH1* **(Fig. 2b).** There are a number of possible explanations for this, which are not mutually exclusive. Firstly, it could simply be a result of a variable background mutation rate across the gene; that is to say, some regions of the gene are inherently more prone to mutation. Synonymous mutations have been used to estimate background mutation rates (43) as they are assumed to be neutrally selected. We see there is a significantly higher number of synonymous mutations in the EGF11-12 region **(Fig. 7a,** p=0.001, n=31, two-tailed Monte Carlo test, **methods)** than the rest of the gene, suggesting that a variable mutation rate could be contributing to the uneven distribution of missense mutations. However, the numbers are low, contain two cases of duplicate mutations within the same donor which could potentially belong to the same clone. It is also possible that some synonymous mutations are not functionally neutral (44) and therefore may be increased/decreased in number due to positive/negative selection, meaning they may not give a true indication of mutation rate. Secondly, it may be that the DNA sequence of the highly mutated regions has a higher probability of acquiring mutations due to the mutational spectrum. However, this does not appear to be the case for missense mutations (Fig. 7b). Thirdly, it could be that a higher proportion of potential mutations are functionally impactful in the ligand-binding region than other sections of the gene. For example, we have seen that approximately two-thirds of potential single nucleotide substitutions in the EGF11-12 repeats are likely to be selected. If this proportion is lower in the rest of the gene, this would result in a smaller number of observed mutations there. Fourthly, the effects of a disruptive mutation may be milder in the non-ligand-binding regions. It might be that disruption of ligand binding completely stops the *NOTCH1* signal, but, for example, a mutation in the transmembrane helix reduces rather than stops the signal (32). Lastly, it has been suggested that Notch receptors may form homodimers or clusters when they bind to a ligand (45, 46). Potentially, this could mean that a mutant ligand-binding region could disrupt the activity of the WT allele as well as the mutant allele, reducing the *NOTCH1* signal more than having completely lost one *NOTCH1* allele (Fig. 5c). This dominant negative effect has been observed for some missense mutations in the *TP53* DNA binding domain (47). The last two hypotheses suggest that the fitness of *NOTCH1* mutant clones depends on the dose of *NOTCH1* signalling. This is consistent with the frequent observation of loss-of-heterozygosity copy number variants associated with *NOTCH1* mutations (4), which strongly suggests that losing both wild type alleles of *NOTCH1* provides a stronger growth advantage than losing a single allele.

**Figure 7:**
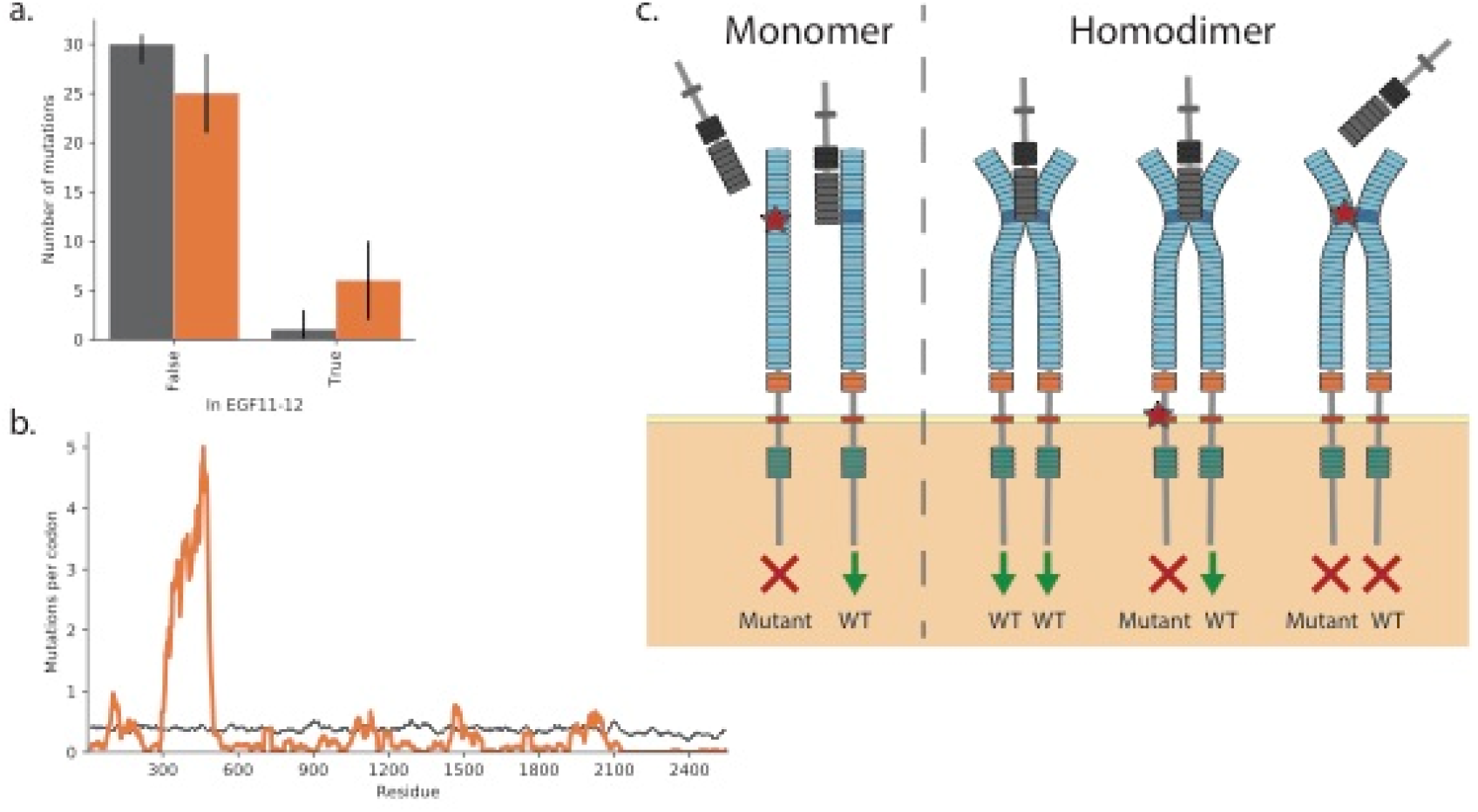
Potential causes of uneven missense mutation distribution across *NOTCH1.* **a.** The counts of synonymous mutations expected under the neutral null model, grey, and observed, orange, in EGF11-12. p=0.001, n=31, two-tailed Monte Carlo test **(methods).** Error bars show 95% confidence intervals **(methods), b.** Sliding window showing the distribution of missense mutations across *NOTCH1* expected under the neutral null model, grey, and observed, orange, **c.** Potential impact of mutations if NOTCH1 (shown with domains as in **Figure 2)** binds to the ligand (grey) as a monomer or a homodimer. If NOTCH1 binds to the ligand as a monomer (left), then a mutation in the ligand-binding EGF repeats may entirely stop signalling from the mutant allele but the wild type (WT) allele still functions normally. If NOTCH1 binds the ligand as a homodimer (right), then a mutant away from the ligand binding region may stop signalling from the mutant allele but not affect the WT allele. However, a mutant in the ligand-binding region may prevent the ligand from binding, and therefore stop signalling from the WT allele as well. If NOTCH1 binds to the ligand in pairs, a heterozygous NOTCH1 mutant would mean only one quarter of NOTCH1 pairs would be WT-WT, so a mutant that prevents binding to the ligand could reduce NOTCH1 signalling by three-quarters.

Some caution must be applied when using the method. Whilst protein misfolding and ligand binding are well known processes controlling protein activity (36, 48), more generally correlation does not mean causation, and significant selection may be found for a feature that correlates (through coincidence or otherwise) with the true selected feature. This is also a test for selection *relative to the rest of the tested region,* and does not necessarily directly translate to positive or negative selection. We have seen that if there are multiple selected features in a region, testing for one individual feature at a time may lead to misleading results, but that this can be corrected by conditioning against the other, confounding features. It also highlights an advantage of using general functional impact scores, rather than focusing on single impact types, when looking for driver genes (17, 18). A future improvement would be to incorporate an estimate for background mutation rate in a region (16, 49) and hence test for absolute rather than relative selection.

Using the statistical method, we can draw out structural and functional information from the increasingly large amount of DNA sequencing data available. Manual investigation of hotspot mutations can provide similar information (4) but can be time consuming, does not leverage the information provided by rarer mutations and does not statistically test the selection of features. The method can be used as an *in vivo* validation of results of *in vitro* studies, or could be a useful method to explore selection of mutation features in existing data sets prior to conducting further experiments. This is an approach that will be widely applicable for genes or domains that are positively or negatively selected in somatic contexts, whether in cancer or normal tissue.

## Methods

### Data

We used mutations detected in normal human oesophagus (4). This experiment used a grid of adjacent oesophagus samples. Large clones could spread over multiple samples. To avoid double counting of such clones, we used the mutations list where mutations that were seen repeatedly in nearby samples were assumed to be from a single clone and were merged (4). The data is available in the original publication (4).

### FoldX

ΔΔG was calculated for every missense mutation in NOTCH1 and NOTCH2 EGF11-12 using FoldX 5 (36) and the pdb files 2VJ3 (37) and 5MWB (28) respectively. First, the FoldX command *RepairPDB* was run to clean up unrealistic residue orientations within the pdb file:

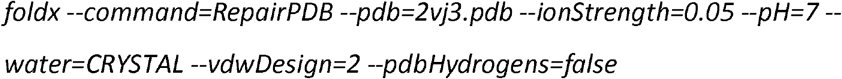

Then the FoldX command *PositionScan* was run for every residue in the structure. This mutates each residue to all other amino acids and calculates the ΔΔG. For example, this command runs for the residue E455:

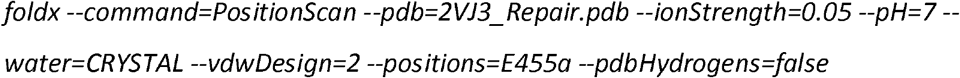

### Ligand-binding interface residues

The ligand-binding residues in EGF11 and EGF12 have been identified for the rat Notch1 bound to the ligands Jag1 and Dll4 (26, 27). Notch genes and ligands are highly conserved between species meaning that the results from the rat protein can be applied to the human NOTCH1 (28). The ligand binding surface is very similar for both ligands (27) and we therefore choose to use the union of both sets of ligand-binding residues **(Supplementary Fig. 1a, Supplementary Table 1).** The ligand binding residues for NOTCH2 are similarly based on conservation with the rat Notch1 ligand interface (28) **(Supplementary Table 2).**

The residue numbers in the pdb files were checked against the residue numbers in the Uniprot protein sequences using SIFTS (50). Distances from the interface residues were calculated using the Python package MDAnalysis (51) using the 2VJ3 structure of NOTCH1 EGF11-13 (37) and using the 5MWB structure of NOTCH2 EGF11-13 (28). They were calculated as the distance from the α-carbon of the residue to the α-carbon of the nearest ligand-interface NOTCH residue. Where alternative locations for an α-carbon existed in the pdb file, the average distance of the locations was used.

### Calcium binding mutations

The calcium binding residues in EGF11-12 of NOTCH1 and NOTCH2 were defined using MetalPDB (38) and the 2VJ3 (37) and 5MWB (28) structures respectively.

### APBS

APBS was run on mutant and wild type structures of the Notch1 structure 2vj3. The mutant structure was generated using the mutate model method in modeller (52). The wild type and mutants structures were relaxed using the Foldx RepairPDB method as follows:

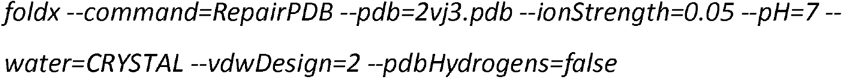

PQR files were generated for APBS calculations using pdb2pqr (53) command line tool and the parse forcefield as follows:

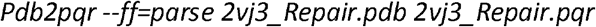

APBS calculations were performed on the resultant PQR file using VMD, with a local installation of the APBS software (53). Electrostatics were calculated with a solute dielectric constant of 8.0 F/m, and solvent dielectric constant of 80.0 F/m. Visualization was performed with VMD (54), using the isosurface representation.

### Conservation scores

Phylop conservation scores (55) for each position were downloaded via the UCSC genome browser, http://hgdownload.soe.ucsc.edu/goldenPath/hg19/phyloP100way/ (56).

### Mutational spectrum calculation

Firstly, all genes containing an exonic mutation in the human oesophagus data set (4) were found using exon locations in GRCh37.p13 downloaded from Ensembl Biomart (57). For each of these genes, the longest transcript was selected, and alternative transcripts discarded. A trinucleotide context was calculated in the direction of the protein transcription for every nucleotide in each transcript and applied to each observed mutation. A mutation rate was calculated for each single nucleotide substitution type in each trinucleotide context by dividing the total number of observations by the number of times the context occurs in the included transcripts.

The mutational spectrum can be assumed to be gene specific (18), to depend on a larger nucleotide context (16), to be distorted by selection hotspot mutations (58), or to be symmetrical in terms of the transcribed strand direction (e.g. AAA>ACA is assumed to have the same mutation rate as to TTT>TGT, so there are 96 possible trinucleotide changes instead of 192) (17). The above method can therefore be adapted by changing the number of context bases, whether to test using a global spectrum for the dataset or one calculated from just the transcript to test, deduplicating recurrent (and therefore possibly selected) mutations before calculating the spectrum, or by assuming there is no difference between mutation rate for *+/-* strands. We show the results of changing these assumptions in **Supplementary Text – Mutational Spectrum.**

Except where otherwise stated, all calculations were made using a trinucleotide, transcribed strand non-symmetric (192) spectrum calculated from all genes in the oesophagus mutation data (4) **(methods – Data).**

### Monte Carlo test

Each possible single nucleotide change was enumerated for the region to be tested. For each of these potential mutations, a relative mutation rate **(methods – Mutational spectrum calculation)** and a metric score (ΔΔG or a Boolean value for whether a mutation was in an interface or calcium binding residue) was calculated. The null hypothesis of neutral selection assumed that the probability of observing each mutation, and therefore the distribution of metric scores, is determined solely by the mutational spectrum. A cumulative distribution of scores under the null hypothesis was calculated, the score for each possible mutation converted to a CDF score, and the sum of these CDF scores taken as the test statistic. Using the summed CDF score instead of the mean of the raw scores for the test statistic means that the test is less sensitive to outlier values. Using the median would be less sensitive to outliers than the mean, but would be inappropriate for testing discrete metrics such as on/off interface. The best choice of statistic to use in the test may depend on the particular data and the question being asked (for example, if the outlier mutations are crucial and reliably scored, then mean may be more appropriate). However, the CDF score sum is a robust option that works for the metric scores tested in this study, and the p-values from the *NOTCH1* Monte Carlo tests using the distribution mean and median are shown in **Supplementary Table 3.**

Let *n* be the number of observed mutations in the region to test. A large number *N* (here N=100000) of random draws of *n* values from null distribution of CDF scores was made, and for each draw, *i,* we calculated the sum of the CDF scores, *s_i_*. The sum of the CDF scores for the observed mutations, *s_obs_* was calculated. We then calculated, *b,* the number of the summed CDF scores *s_i_* that are more extreme than the observed summed CDF score. This was converted into a two-tailed p-value by multiplying the exact Monte Carlo p-value (59) by two, i.e.

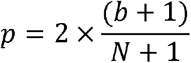

where

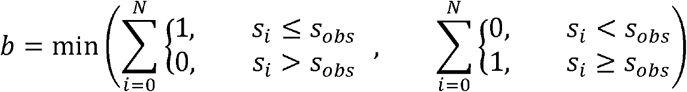

The minimum p-value possible with N=100000 is just under 2e^-5^ (2/100001). Except where stated otherwise, p-values in this manuscript use this test.

### Confidence intervals

95% confidence intervals were calculated by taking 10000 random samples from the null distribution or by bootstrapping 10000 random samples with replacement from the observed data.

## Acknowledgements

We thank the Hall and Jones groups for discussions. This work has been supported by the Royal Society (URF to BAH grant no. UF130039), Medical Research Council (Grant-in-Aid to the MRC Cancer unit and NIRG to BAH, grant no. MR/S000216/1), the Harrison Watson Fund at Clare College, Cambridge (M.W.J.H.), Cancer Research UK Programme (P.H.J., grant no. C609/A17257), & the Wellcome Trust (to the Wellcome Sanger Institute, 098051 and 206194).

---

**Supplementary Figure 1:**
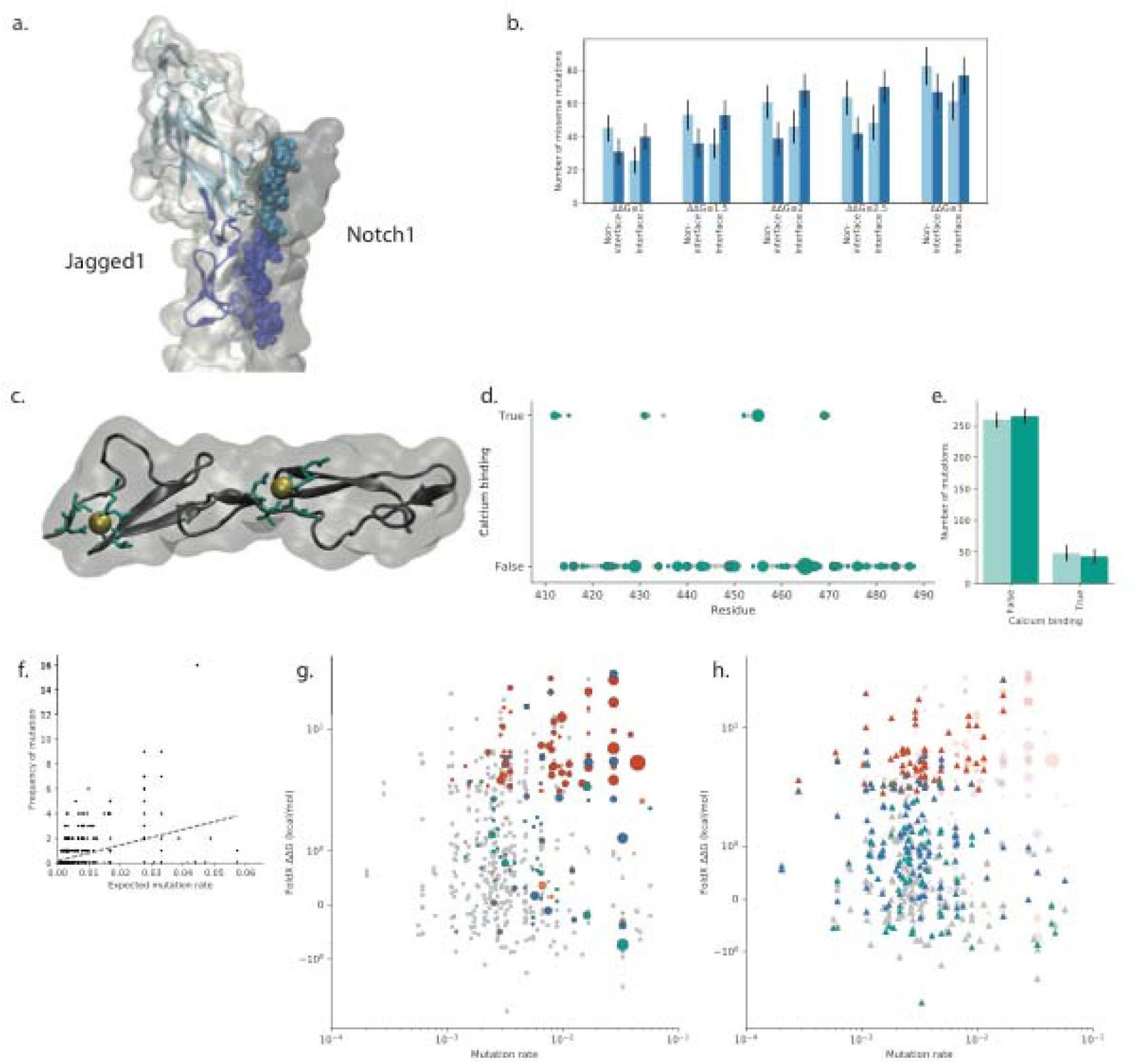
Missense mutations in *NOTCH1* EGF11-12. **a.** Ligand interface residues **(methods)** shown on rat Notch1 bound to Jagged1 (PDB 5UK5). Site 1 interface residues in Notch1 EGF12 and the Jagged1 C2 domain shown in light blue, Site 2 interface residues in Notch1 EGF11 and the Jagged1 DSL domain shown in dark blue. **b.** Number of EGF11-12 mutations in/not in ligand-binding interface residues when excluding mutations which are highly destabilising. Expected counts under the “neutral” null hypothesis show in light blue, observed counts in dark blue. The ΔΔG thresholds for excluding destabilising mutations are 1, 1.5, 2, 2.5, 3 kcal/mol. For each of the thresholds, there is a significant enrichment of observed interface mutations (p-values all < 0.0002, two-tailed Monte Carlo test, **methods).** Error bars show 95% confidence intervals **(methods), c.** Calcium-binding residues in *NOTCH1* EGF11-12 **(methods).** Calcium-binding residues shown in green, Ca^2+^ ions shown in yellow, **d.** Mutations on calcium-binding residues in *NOTCH1* EGF11-12. Single nucleotide missense mutations that occur in the human oesophagus data set, green, with marker size proportional to the number of times that mutation recurs. Single nucleotide missense mutations that do not occur in the human oesophagus data set shown in grey. **e.** Counts of *NOTCH1* EGF11-12 mutations that are on calcium-binding residues, under the neutral null hypothesis, light green, and observed, dark green. p=0.48, two-tailed Monte Carlo test **(methods).** Error bars show **95%** confidence intervals **(methods), f.** Expected mutation rate **(methods)** vs number of observations for each potential mutation in *NOTCH1* EGF11-12. There is a trend of increasing number of occurrences with higher expected mutation rate (R^2^=0.16; p-value=4e^-21^, two-tailed Wald Test for zero slope using scipy stats.linregress (61)). **g.** FoldX ΔΔG plotted against relative expected mutation rate for missense mutations in *NOTCH1* EGF11-12. Single nucleotide missense mutations that occur in the human oesophagus data set with marker size proportional to the number of times that mutation occurs shown in green if the residue is calcium binding, blue if the residue is on the ligand-binding interface, red if the mutation has ΔΔG>2kcal/mol, orange otherwise. Single nucleotide missense mutations that do not occur in the human oesophagus data set shown in grey. **h.** FoldX ΔΔG plotted against relative expected mutation rate for missense mutations in *NOTCH1* EGF11-12. Unobserved mutations shown as triangle, green if the residue is calcium binding, blue if the residue is on the ligand binding interface, red if the mutation has ΔΔG>2kcal/mol, grey otherwise. Observed mutations shown as faded circles with marker size proportional to the number of times that mutation occurs and colours as in **g.** There are potential mutations that are highly destabilising, on the ligand-binding interface or calcium binding and with relatively high expected mutation rates that are not observed in the normal human oesophagus data set.

**Supplementary Figure 2:**
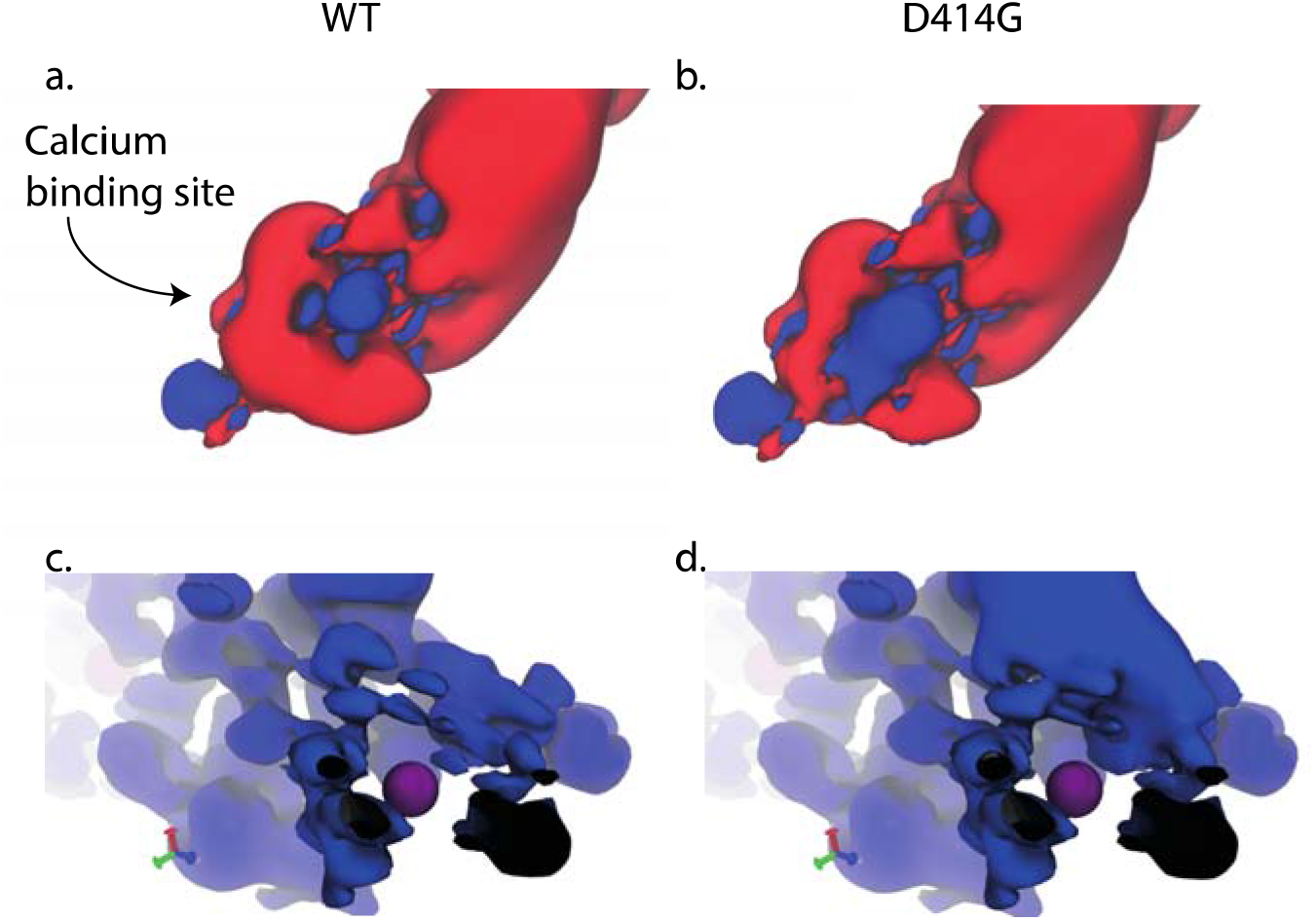
Adaptive Poisson-Boltzmarm Solver calculations for NOTCH1 2vj3. Shown are A) Isosurface for wild type (WT) NOTCH1where blue represents areas of positive charge, and red areas of negative charge. B) Isosurface for D414G mutant NOTCH1. Also shown is positive charge only around the D414 adjacent calcium ion for: C) WT, and D) D414G mutant structures.

**Supplementary Figure 3:**
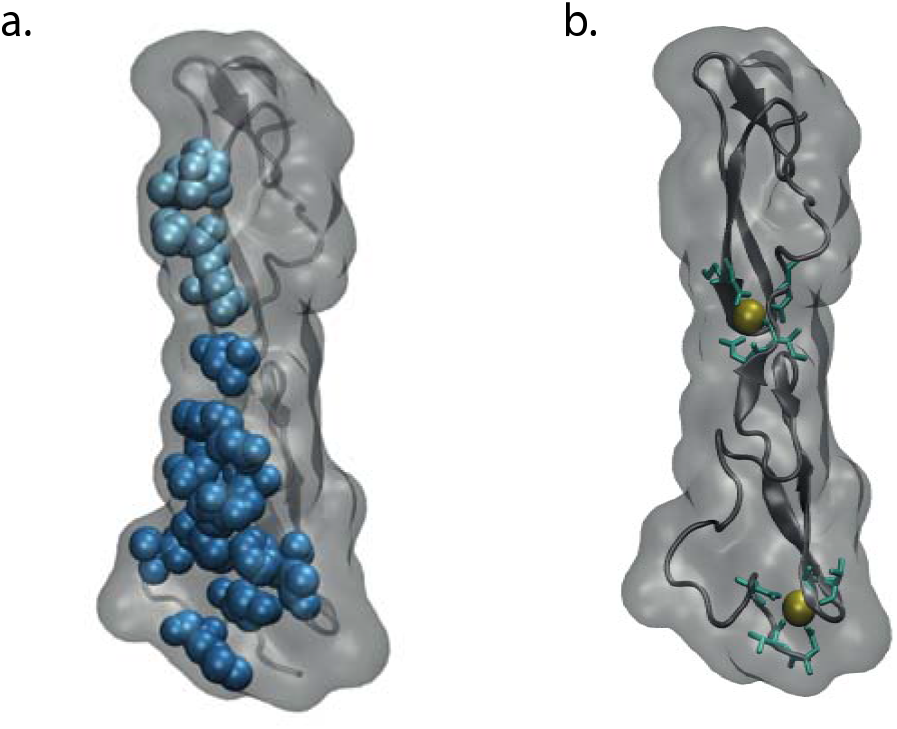
Interface and calcium binding residues in *N0TCH2* EGF11-12. **a.** Ligand interface residues **(methods)** of (PDB 5UK5). Site 1 interface residues in *NOTCH2* EGF12 shown in light blue, Site 2 interface residues in *NOTCH2* EGF11 shown in dark blue. **b.** Calcium-binding residues in *NOTCH2* EGF11-12 **(methods).** Calcium-binding residues shown in green, Ca^2+^ ions shown in yellow.

**Supplementary Figure 4:**
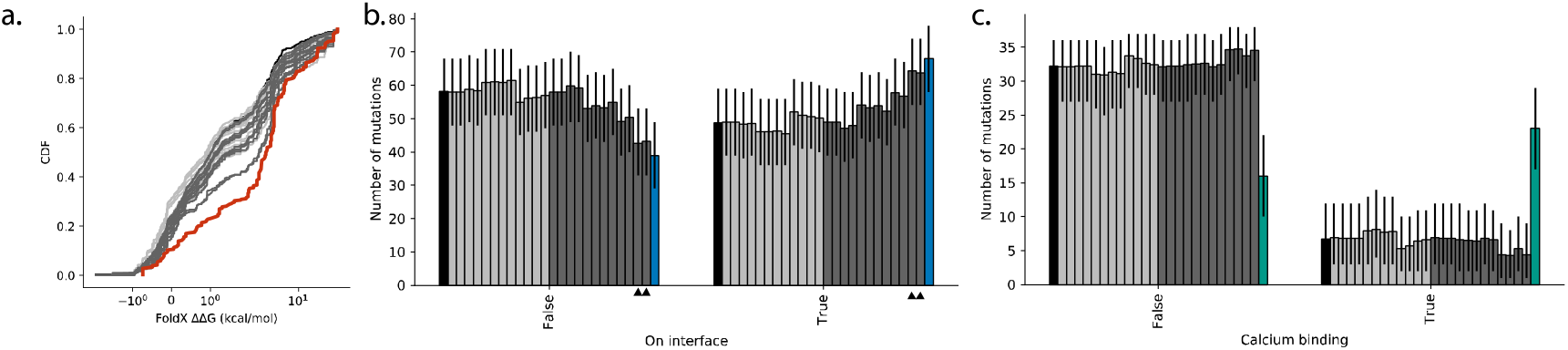
Mutational spectrum assumptions, **a.** Cumulative distributions of the observed ΔΔG values, red, and the null distributions, grey and black. The observed distribution is significantly different from all shown null distributions (p<0.002, n=308, twotailed Monte Carlo test, **methods).** Null distribution using a spectrum with all mutations with equal probability shown in black, null distributions with spectra calculated from mutations in all genes in the data set shown in light grey, and null distributions with spectra calculated only from mutations in *NOTCH1* shown in dark grey. Full list of p-values shown in **Supplementary Table 4. b.** Counts of *NOTCH1* EGF11-12 mutations with calculated ΔΔG ≤ 2 kcal/mol occurring on the ligand-binding interface under the null hypotheses, grey or black, and observed, blue. Colours of null hypotheses as in **a.** Error bars show 95% confidence intervals **(methods).** The observed distribution is significantly different from all but two of the shown null distributions (p<0.02 for these significant cases, n=107, two-tailed Monte Carlo test, **methods).** The exceptions are the pentanucleotide, transcribed/non-transcribed strand separating, per-gene spectra – indicated with arrowheads. Full list of p-values shown in **Supplementary Table 4. c.** Counts of *NOTCH1* EGF11-12 mutations that are on calcium-binding residues (having excluding mutations with calculated ΔΔG > 2 kcal/mol or occurring on the ligand-binding interface), under the null hypotheses, grey or black, and observed, green. Colours of null hypotheses as in **a.** The observed distribution is significantly different from all shown null distributions (p<2e^-5^, n=39, two-tailed Monte Carlo test, **methods).** Error bars show 95% confidence intervals **(methods).** Full list of p-values shown in **Supplementary Table 4.**

**Supplementary Figure 5:**
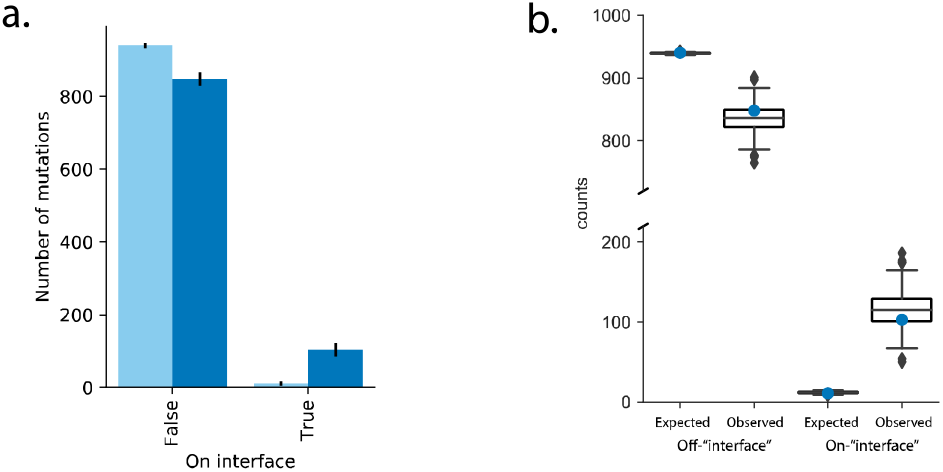
Altering the test region, **a.** Counts of NOTCH1 mutations occurring on the EGF11-12 ligand-binding interface under the neutral null hypothesis, light blue, and observed, dark blue. p<2e^-5^, n=951, two-tailed Monte Carlo test **(methods).** Error bars show 95% confidence intervals **(methods), b.** Boxplots showing the expected and observed counts of NOTCH1 mutations observed in 1000 random subsets of 29 EGF11-12 residues (same number as in the EGF11-12 ligand-binding interface). Counts for the EGF11-12 ligand-binding interface shown as blue dots. In all cases, there is significant selection of mutations in the subset of EGF11-12 residues (p<2e^-5^ in all cases, n=951, two-tailed Monte Carlo test, **methods**).

**Supplementary Figure 6:**
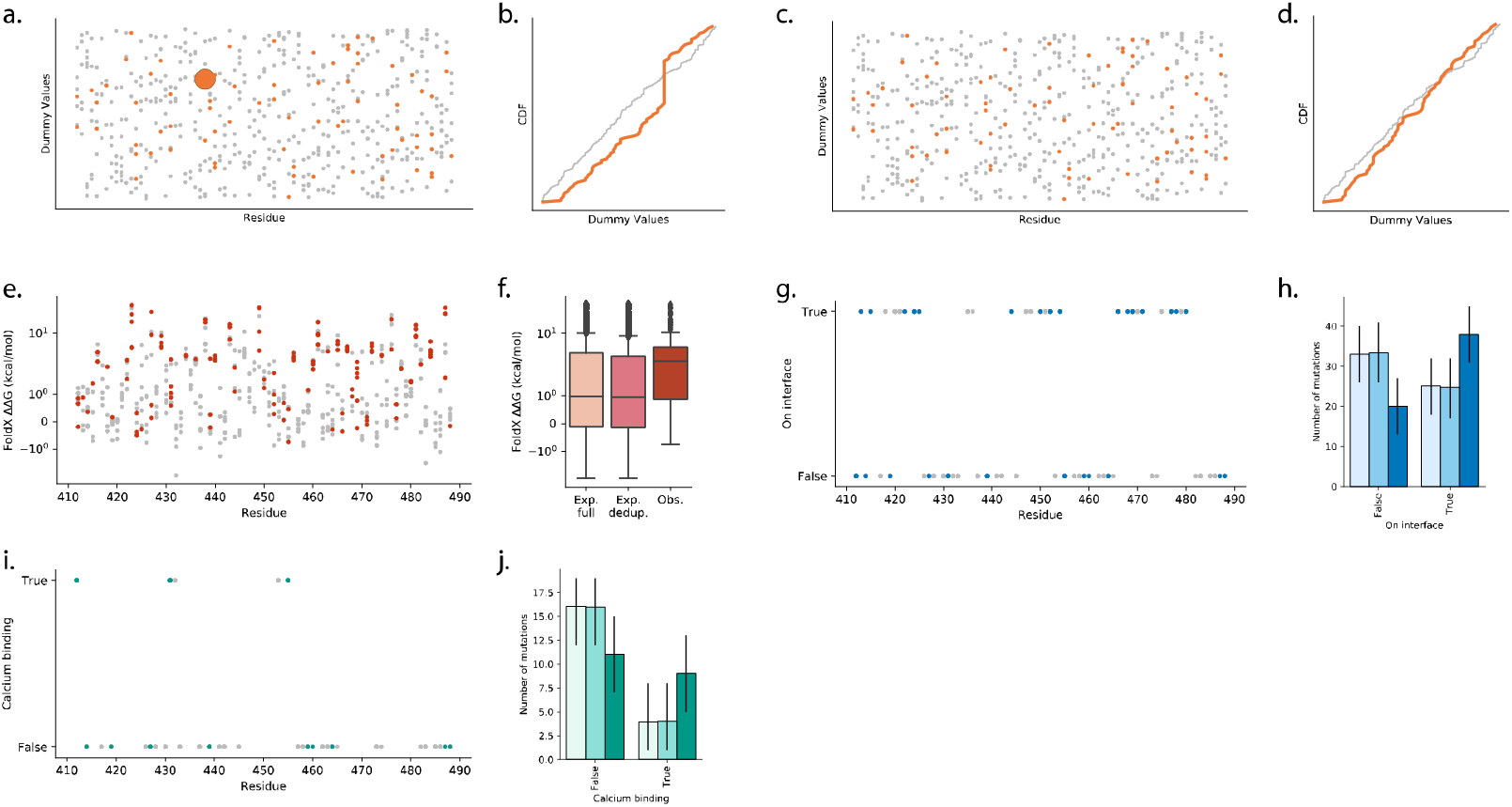
The effect of hotspot mutations, **a-d,** demonstration of how a hotspot mutation can lead to rejection of the null hypothesis even when the metric has no biological meaning. A synthetic set of mutations was generated, where 74 mutations appear once, and one hotspot mutation appears 25 times. In place of a metric based on a biological feature, random scores between zero and one are generated for each mutation, **a.** Scatter plot of the example mutations. “Observed” mutation shown in orange, “unobserved” mutations in the region shown in grey. The hotspot mutation is the large orange marker, **b.** Cumulative distributions of the metric for the null model, grey, and the “observed” mutations. The large vertical jump in the “observed” CDF is caused by the hotspot mutation. The “observed” distribution is significantly different from the null (p=0.005, two-tailed Monte Carlo test, **methods), c.** Scatter plot of the example mutations after removing duplicate mutations. “Observed” mutation shown in orange, “unobserved” mutations in the region shown in grey. The hotspot mutation is now only counted once. **d.** Cumulative distributions of the metric for the null model, grey, and the “observed” mutations after removing duplicate mutations (p=0.3, two-tailed Monte Carlo test, **methods), e-j,** reanalysis of the selection of features in *NOTCH1* EGF11-12 after removing duplicate mutations, **f, h, j** two null models are tested, either using the full set of mutations to calculate the mutation spectrum (lightest shade, leftmost bar/boxplot) or deduplicating the mutations prior to calculating the spectrum (middle shade, middle bar/boxplot). Observed mutations shown in the darkest shade, rightmost bar/boxplot. **e.** ΔΔG of mutations in *NOTCH1* EGF11-12 calculated using structure 2VJ3 and FoldX. Single nucleotide missense mutations that occur in the human oesophagus data set, red. Single nucleotide missense mutations that do not occur in the human oesophagus data set shown in grey. **f.** Distribution of calculated ΔΔG values of missense mutations. Distribution expected under the neutral null hypothesis using all mutations for the spectrum, left, distribution expected under the neutral null hypothesis using deduplicated mutations for the spectrum, middle, and the distribution observed after removing duplicate mutations, right. p<2e-5 for both null models, n=139, two-tailed Monte Carlo test **(methods), g.** Mutations on or off the ligand-binding interface of *NOTCH 1* EGF11-12 **(methods),** where mutations with a calculated ΔΔG>2 kcal/mol have been excluded. Single nucleotide missense mutations that occur in the human oesophagus data set, blue. Single nucleotide missense mutations that do not occur in the human oesophagus data set expected to occur on the ligand-binding interface. Counts under the neutral null hypothesis using all mutations for the spectrum, lightest blue (p=0.001), under the neutral null hypothesis using deduplicated mutations for the spectrum, mid-blue (p=0.0008), and observed after removing duplicate mutations, dark blue. n=58, two-tailed Monte Carlo test **(methods).** Error bars show 95% confidence intervals **(methods).** ¡. Mutations on calcium-binding residues in *NOTCH1* EGF11-12 where mutations with a calculated ΔΔG>2 kcal/mol or on a ligand-binding interface residue have been excluded. Single nucleotide missense mutations that occur in the human oesophagus data set, green. Single nucleotide missense mutations that do not occur in the human oesophagus data set shown in grey. **j.** Counts of *NOTCH1* EGF11-12 mutations that are on calcium-binding residues (having excluding mutations with calculated ΔΔG > 2 kcal/mol or occurring on the ligand-binding interface). Counts under the neutral null hypothesis using all mutations for the spectrum, lightest green, under the neutral null hypothesis using deduplicated mutations for the spectrum, mid-green, and observed after removing duplicate mutations, dark green. p=0.02 for both null models, n=20, two-tailed Monte Carlo test **(methods).** Error bars show 95% confidence intervals **(methods).**

## Supplementary Text

In this supplementary text we explore the assumptions of the statistical method, discuss potential limitations and describe methods that can be used to test the robustness of results.

### Mutational spectrum

Cells acquire somatic mutations through exposure to mutagens or during (sometimes defective) cellular processes (13). Different mutational processes have their own characteristic ‘signatures’ of nucleotide substitution frequencies that depend on the nucleotide change itself and the surrounding nucleotide context (13). The pattern of mutations observed in a sample, known as the mutational spectrum, is a mix of the signatures of the contributing mutational processes (13).

The probability of a particular somatic mutation appearing in a sequenced sample depends on both the rate at which the mutation occurs in cells and the strength of selection on the mutation once it has occurred. We want to separate these two factors, and therefore require a model of how often each mutation would appear in the absence of selection (the null hypothesis model). A common method to do this is to assume that, within a gene, each mutation in the same spectrum category has the same mutation rate (16–18). For example, using a trinucleotide spectrum, two AAA trinucleotide sequences at different locations within the same gene are assumed to have the same probability of mutating into ACA.

A variety of spectrum assumptions have been used in driver detection models. Using a trinucleotide mutational spectrum to generate a null model has been found to improve driver detection compared to more simplistic substitution models (16). However, it has also been found that a pentanucleotide context is more appropriate for the signature of ultraviolet light (16). Transcription coupled repair may produce a strand bias in the mutational spectrum (16) which can be taken into account by separating the transcribed and non-transcribed strand of a gene. Previous studies have assumed either that all coding regions in the genome share the same mutational spectrum (16, 17), or that each gene has its own spectrum (18).

Each of these assumptions alters the number of mutation rate parameters we need to estimate. The more mutation rate parameters required, the more thinly spread the observed mutations, and the noisier the estimation of each rate will become. The choice of assumptions is therefore a compromise between a potentially more accurate representation of the context that determines mutation rate (larger spectrum) and more stable estimates of each rate parameter (smaller spectrum).

Selected mutations within the data may distort the mutational spectrum. This could be a larger problem when using gene specific spectra or for data sets containing only a small number of genes because the selected mutations may make up a larger proportion of the total data. To some extent this selection bias can be reduced by removing duplicate mutations before calculating the spectrum (58). For high mutation loads, where duplicate mutations are likely even under neutral selection, this deduplication may have its own distorting effect.

We have run the tests for *NOTCH1* EGF11-12 (the same tests as in the **EGF repeat mutations cause misfolding and disrupt ligand and calcium binding sites** section of **Results)** using a range of different spectrum assumptions to see how robust the signs of selection are to a variety of null hypotheses. The selection of destabilising mutations is highly significant for all of the tested spectra (p<0.002, **Supplementary Fig. 4a, Supplementary Table 4),** as is the selection of mutations on calcium binding residues (p< 2e^-5^, two-tailed Monte Carlo test **-methods,** excluding interface and destabilising mutations – see main text, **Supplementary Fig. 4c).** The selection of mutations on the ligand-binding interface is significant for all spectra apart from the pentanucleotide, transcribed/non-transcribed strand separating, per-gene spectra (p>0.4, p<0.05 for all other spectra, two-tailed Monte Carlo test – **methods,** excluding destabilising mutations – see main text, **Supplementary Fig. 4b).** This is an example of an overly detailed spectra for the data (18), as it has 3072 separate rate parameters, calculated from less than 1300 single nucleotide substitutions in NOTCH1.

We may be able to further improve the null models in future as work continues to determine the factors that influence mutation rate (62)

### Testing region

Typically, when searching for selection in somatic mutations, the mutations are grouped by gene (16–18). However, we are not just interested in which genes are under selection, but which mutations in the gene are selected and why. The method **(Fig. 1)** looks at relative selection within a defined region, effectively testing whether the subset of mutations that share a certain feature are enriched *compared to the rest of the mutations in the region.* Therefore, not only the choice of the feature, but the choice of the region can affect the results.

The region definition is not limited to contiguous sections of protein sequence, but the choice of which mutations within those sequences to include in the test. Whenever certain mutations are excluded (whether by location or a metric value), they must be excluded from both the null model and the observed mutations to avoid biasing the test.

We will use the EGF11-12 interface mutations to demonstrate the impact the choice of test region can have on the results. Using the whole of the NOTCH1 protein as the test region, we found that there is a significant increase of missense mutations on the EGF11-12 ligand-binding interface residues (p<2e^-5^, n=951, two-tailed Monte Carlo test – **methods, Supplementary Fig. 5a).** However, the rest of the gene has a far lower density of mutations than EGF11-12 (Fig 2b, see **Discussion).** Mutations on randomly selected groups of residues in EGF11-12 were also significantly increased (1000 random groups of N residues in EGF11-12 tested, p<2e^-5^ in each case, n=951, two-tailed Monte Carlo test – **methods, Supplementary Fig. 5b).** Therefore, using the whole gene as the test region, we cannot show if the interface residues are under particularly strong selection compared to an arbitrary group of residues in EGF11-12.

Using EGF11-12 as the test region, we have seen that there is no significant difference in selection on mutations on the interface compared to the all other mutations in EGF11-12 **(Fig. 3c,d, Results),** but *non-destabilising* mutations were significantly more likely to occur on the ligand-binding interface **(Fig. 3e,f, Results).** We conclude that the interface mutations in EGF11-12 are particularly strongly selected, and therefore are (unsurprisingly) important for the binding of NOTCH1 to its ligand.

The choice of region is a compromise. Larger regions increase the sample size, meaning more power to detect significant results. However, the assumptions used in the null model (such as constant mutation rate and spectrum) may be less likely to hold over large distances. When using metrics which require a structure to calculate, such as ΔΔG, it may necessitate running over a small region. And we have seen that when faced with multiple selected features, it may be necessary to test one feature while exclude mutations using the others.

As our example makes clear, altering the test region changes the question being asked by the test: *Are the interface mutations more selected than the bulk of mutations in NOTCH1? Are the interface mutations more selected than the bulk of mutations in EGF11-12? Are the non-destabilising interface mutations more selected than the bulk of the non-destabilising mutations in EGF11-12?* The user must decide which question(s) are most relevant for their interest.

### Correlation, causation and hotspots

The statistical method tests for a shift in the distribution of a feature metric between the null model and the observed mutations **(Fig. 1).** Ideally, any shift detected will be due to selection of the particular feature that is being tested. However, hotspots can distort the distribution, regardless of the metric used, and can even provide evidence of selection when using random numbers for the metric (18). This is not necessarily a problem when just looking for evidence of selection acting on a gene. However, we are interested in whether the particular feature is selected, and distortions due to hotspots may give a false impression of the importance of a feature **(Supplementary Fig. 6a,b)**

A simple approach is to check that the trend of selection is still apparent after deduplicating the recurrent mutations **(Supplementary Fig. 6c).** P-values from this approach may not be reliable because hotspots may be expected to occur if the feature is strongly selected, and deduplicating mutations both reduces the sample size and could distort the mutational spectrum used for the null model **(Supplementary Text – Mutational Spectrum).** However, it can still be a useful method to explore the influence of hotspots.

We used two null models: one with a spectrum calculated from the full data, as used in the main figures; the other with a spectrum calculated using deduplicated data **(Supplementary Text – Mutational Spectrum).** We see that the trends of selection in *NOTCH1* EGF11-12 for destabilising mutations, ligand-binding interface mutations and calcium-binding mutations are still apparent after the removal of duplicate mutations **(Supplementary Fig. 6).** This gives us more confidence that the results shown in the Results section are due to the selection of the features we have identified.

**Supplementary Table 1:**
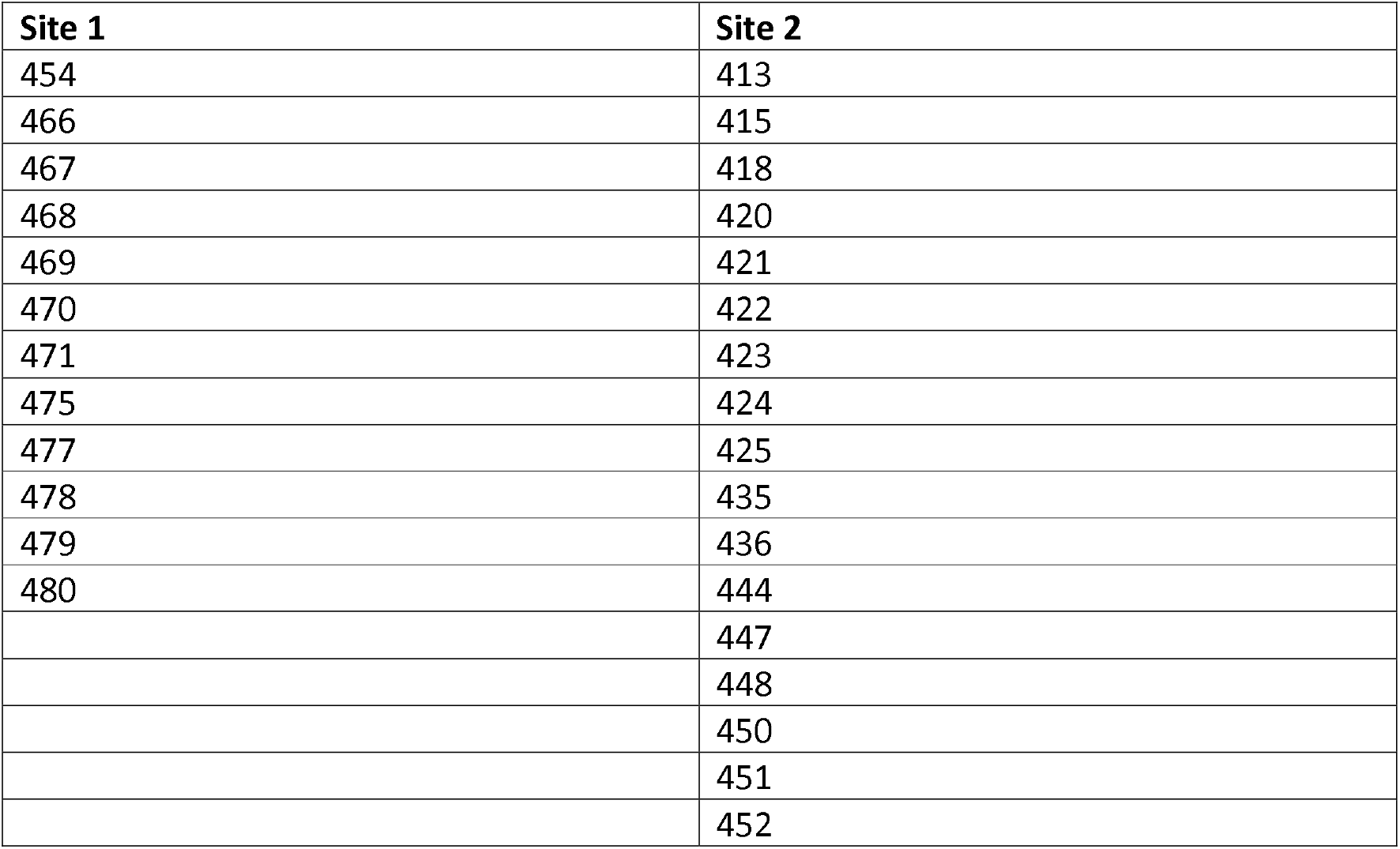
Ligand binding interface residues of NOTCH1 EGF11-12. These residues are taken from Figure S3 of (27) and split into Site 1 (C2 binding) and Site2 (DSL binding) based on (26).

**Supplementary Table 2:** Ligand binding interface residues of NOTCH2 EGF11-12. These residues are taken from (28).

| Site 1 | Site 2 |
| --- | --- |
| 470 | 418 |
| 472 | 421 |
| 473 | 424 |
| 481 | 425 |
|  | 426 |
|  | 428 |
|  | 429 |
|  | 439 |
|  | 440 |
|  | 452 |
|  | 454 |
|  | 456 |

**Supplementary Table 3:** Monte Carlo test p-values using summed CDF score, mean or median as the distribution statistic, see **methods.**

| Metric | Data | Monte Carlo Test P-values |  |  |
| --- | --- | --- | --- | --- |
|  |  | CDF score | Mean | Median |
| FoldX $\Delta\Delta G$ | All missense | $2e^{-5}$ | $2e^{-5}$ | $2e^{-5}$ |
| Ligand-binding interface (bool) | All missense | 0.73 | 0.73 | 1 |
| Ligand-binding interface (bool) | Missense excluding $\Delta\Delta G > 2\text{kcal/mol}$ | $4e^{-5}$ | $4e^{-5}$ | 0.15 |
| Calcium-binding residues (bool) | All missense | 0.48 | 0.48 | 1 |
| Calcium-binding residue (bool) | Missense excluding $\Delta\Delta G > 2\text{kcal/mol}$ | $2e^{-5}$ | $2e^{-5}$ | $2e^{-5}$ |

**Supplementary Table 4:**
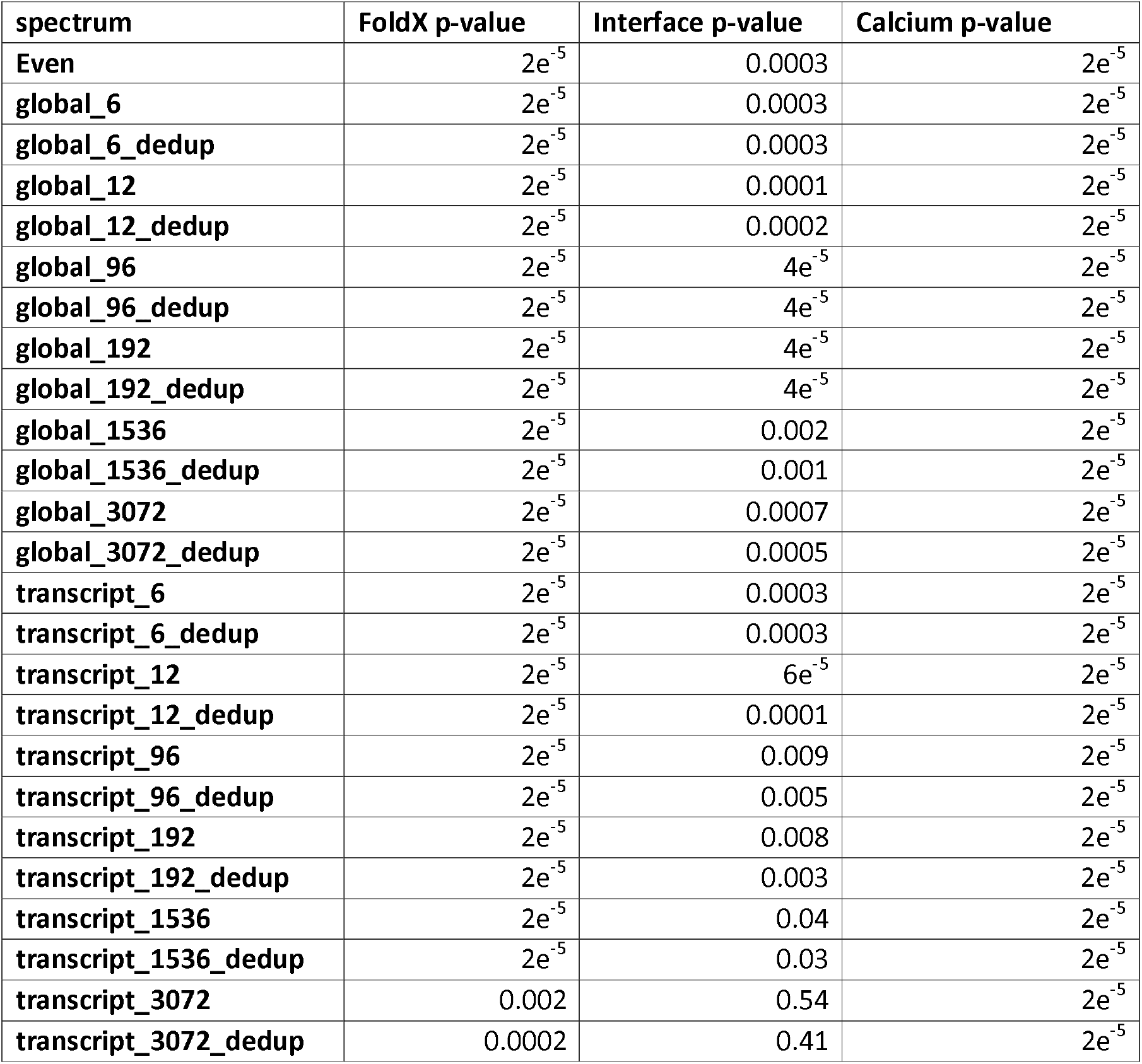
Monte Carlo test p-values under null models that use different assumptions for the mutational spectrum. These are the equivalent tests to those in **Figure 3b,f,î.** The “Even” spectrum assigns an equal probability to every mutation. The “global” spectra calculate the mutation rates using all exonic single nucleotide mutations in the data set. The “transcript” spectra calculate the mutations rates using only the exonic single nucleotide mutations in *NOTCH1.* The number in the spectrum name is the number of mutation rates in the spectrum. “6” and “12” do not use a wider nucleotide context, “96” and “192” use a trinucleotide context, and “1536” and “3072” use pentanucleotide contexts.”6”, “96” and “1536” do not distinguish between the transcribed and nontranscribed strands, “12”, “192” and “3072” do. “dedup” indicates that duplicate mutations (defined as having the same chromosomal position, reference base and mutant base) are removed before calculating the spectrum. The lowest possible p-value from the Monte Carlo tests used was 2e^-5^. The global_192 spectrum has been used throughout this study unless otherwise specified.

